# PopInf: An approach for reproducibly visualizing and assigning population affiliation in genomic samples of uncertain origin

**DOI:** 10.1101/823344

**Authors:** Angela M. Taravella Oill, Anagha J. Deshpande, Heini M. Natri, Melissa A. Wilson

## Abstract

Germline genetic variation contributes to cancer etiology, but self-reported race is not always consistent with genetic ancestry, and samples may not have identifying ancestry information. Here we describe a flexible computational pipeline, PopInf, to visualize principal components analysis output and assign ancestry to samples with unknown genetic ancestry, given a reference population panel of known origins. PopInf is implemented as a reproducible workflow in Snakemake with a tutorial on GitHub. We provide a pre-processed reference population panel that can be quickly and efficiently implemented in cancer genetics studies. We ran PopInf on TCGA liver cancer data and identify discrepancies between reported race and inferred genetic ancestry.

**Significance:** The PopInf pipeline facilitates visualization and identification of genetic ancestry across samples, so that this ancestry can be accounted for in studies of disease risk. All code and a tutorial are available on Github: https://github.com/SexChrLab/PopInf.

## INTRODUCTION

Cancer is a complex disease with genetic and environmental factors contributing to its risk and progression. The underlying genetic architecture of cancer, like other complex diseases, is influenced by common population-specific genetic variation (1,2). Common genetic variation is shared within populations of shared genetic ancestry. Unaccounted population structure can confound the results of genetic analyses, like in cancer GWAS, by causing spurious associations to disease phenotypes (3). Thus, assessing genetic ancestry and population structure in studies on the effects of genetic loci and genetic background on cancer is crucial.

Cancer research has begun to recognize the importance of identifying genetic ancestry across patients in cancer genetic datasets (4) and across cancer cell lines (5). Yuan et al. (4) characterized genetic ancestry across The Cancer Genome Atlas (TCGA) patient cohort to investigate the effect genetic ancestry has on genomic alterations across different cancers and to provide researchers with detailed ancestry information on each patient. Though this publicly accessible resource is of great research value for those using the TCGA data, researchers utilizing other datasets will have to independently infer the ancestry of their samples.

Methods and software are currently available to characterize population structure (6,7), estimate local and global ancestry proportions (7,8), or predict ancestry using genomic data (9). These rely on a pre-defined reference panel and may not report admixed samples. Having an easily reproducible and modifiable workflow to visualize PCA and identify ancestry in individuals of unknown ancestral origin would thus be a useful addition to the cancer genetics researchers tool kit.

Here we present PopInf v1.0, a pipeline to visualize PCA output and assign ancestry to individuals with unknown ancestry, given a flexible reference population panel of known origins. PopInf v1.0, takes, as input, variants from a sample with unknown or unverified genetic ancestry in variant call format (VCF), compares the variants in the unknown sample to a user defined reference panel, and outputs an inferred ancestry origin report with accompanying PCA plots of the unknown samples and the reference panel. We ran PopInf on variants from 148 samples from the Genotype Tissue Expression (GTEx) Project (10) and on 403 samples from the TCGA liver cancer dataset (11) and identify discrepancies between reported race and inferred genetic ancestry. Further, we analyze each sample by chromosome and find cases of chromosome-specific admixture that is not reported in genome-wide analyses.

## MATERIALS AND METHODS

PopInf v1.0 uses a combination of publicly available software and custom scripts to generate PCA plots and a tab-delimited inferred ancestry report for samples of unknown ancestry or unverified self-reported population ancestry. PopInf v1.0 uses GATK v3.7 (12), VCFtools v.0.1.14 (13), bedtools v.2.27.1 (14), and Plink v.1.9 (15) to prepare the unknown ancestry dataset and reference panel, smartpca - a program within EIGENSOFT v6.0.1 package (6) - for PCA, and a custom R script (16) to infer individuals ancestry and plot the results of PCA of the study samples and reference panel. Our pipeline is incorporated into the reproducible workflow system, Snakemake v5.4.0 (17).

### Input

Two sets of variant data are required to use PopInf v1.0: 1) variants from reference populations, and 2) variants from sample(s) of unknown or self-reported race or ancestry. These files need to be mapped to the same reference genome and in VCF file format. Additionally, two sample information text files, one for the reference panel and one for the unknown dataset, are needed for input, each with three tab-delimited columns. For the reference panel sample information text file, column one must contain sample names identical to the naming in the VCF file with one sample per row; column two must specify genetic sex information (“Male” “Female” or “N/A” if unknown, case insensitive); column three must contain population assignment. For the study sample information text file, columns one and two are similar to the reference panel file, but column three is a dummy variable with a single arbitrary value that is the same on every row. For example, column three of the sample information text file for the unknown set of samples could be set as “unknown”. Finally, the user must provide the FASTA file (.fa) of the reference genome used for read mapping along with a FASTA index file (.fai) and a sequence dictionary file (.dict).

### Data processing

PopInf v1.0 implements filtering, merging, and file conversion prior to PCA. Single nucleotide polymorphisms (SNPs) are extracted from both the reference panel and study sample VCF files, using GATK v3.7 SelectVariants and merged using GATK v3.7 CombineVariants (12). To ensure PopInf analyzes SNPs that overlap with both the reference and unknown variant sets, missing genotype data is removed using VCFtools v.0.1.14 (vcftools --max-missing flag) (13). If analyzing the X chromosome, the pseudoautosomal regions and X-transposed region (18,19) are masked using bedtools v.2.27.1 (14). Prior to running PCA, the merged VCF file is pruned for linkage disequilibrium (LD) and converted to plink format using Plink v1.9 (15). PCA on a user-defined set of chromosomes (e.g. whole genome, all autosomes, or a single chromosome) is carried out using smartpca (6).

### Output

PopInf v1.0 generates PCA plots for the first ten PCs for the study samples and the reference panel, and an inferred ancestry report. Genetic ancestry of each study sample is inferred based on the distance between the study sample and the centroid coordinates of PCs 1 and 2 of each reference population. A study sample is inferred to originate from a particular population if it falls within N standard deviations (SDs) from the reference population centroid. To provide multiple levels of confidence, the ancestry is inferred using 1, 2, and 3 SDs. If the sample does not fall within three standard deviations of any population, the sample’s ancestry will be assigned to the closest population or will be assigned as having admixed ancestry: PopInf calculates the midpoint coordinates between each pairwise combination of reference populations and then compares those distances to the study sample. For a sample to be assigned as admixed, it must be closer to the midpoint of two populations than to the 3rd standard deviation of any population. If the study sample is closer to the 3rd standard deviation of a population than any of the midpoints, it will be assigned to that population.

## RESULTS

### Usage examples

We ran PopInf v1.0 using variants from two human genetic datasets: one from healthy individuals and one from cancer patients. The GTEx Project (10) dataset consisted of 148 individuals and the TCGA liver cancer dataset (11) consisted of 403 individuals (Supplementary Table 1; Supplementary Table 2; Figure 1). Both datasets included self-reported race for most individuals. We inferred the genetic ancestries of these samples based on a reference panel consisting of variants from 986 unrelated individuals from populations across Africa, Europe, East Asian, and South Asia from 1000 Genomes Release 3 (20) (Supplementary Table 3).

**Figure 1.**
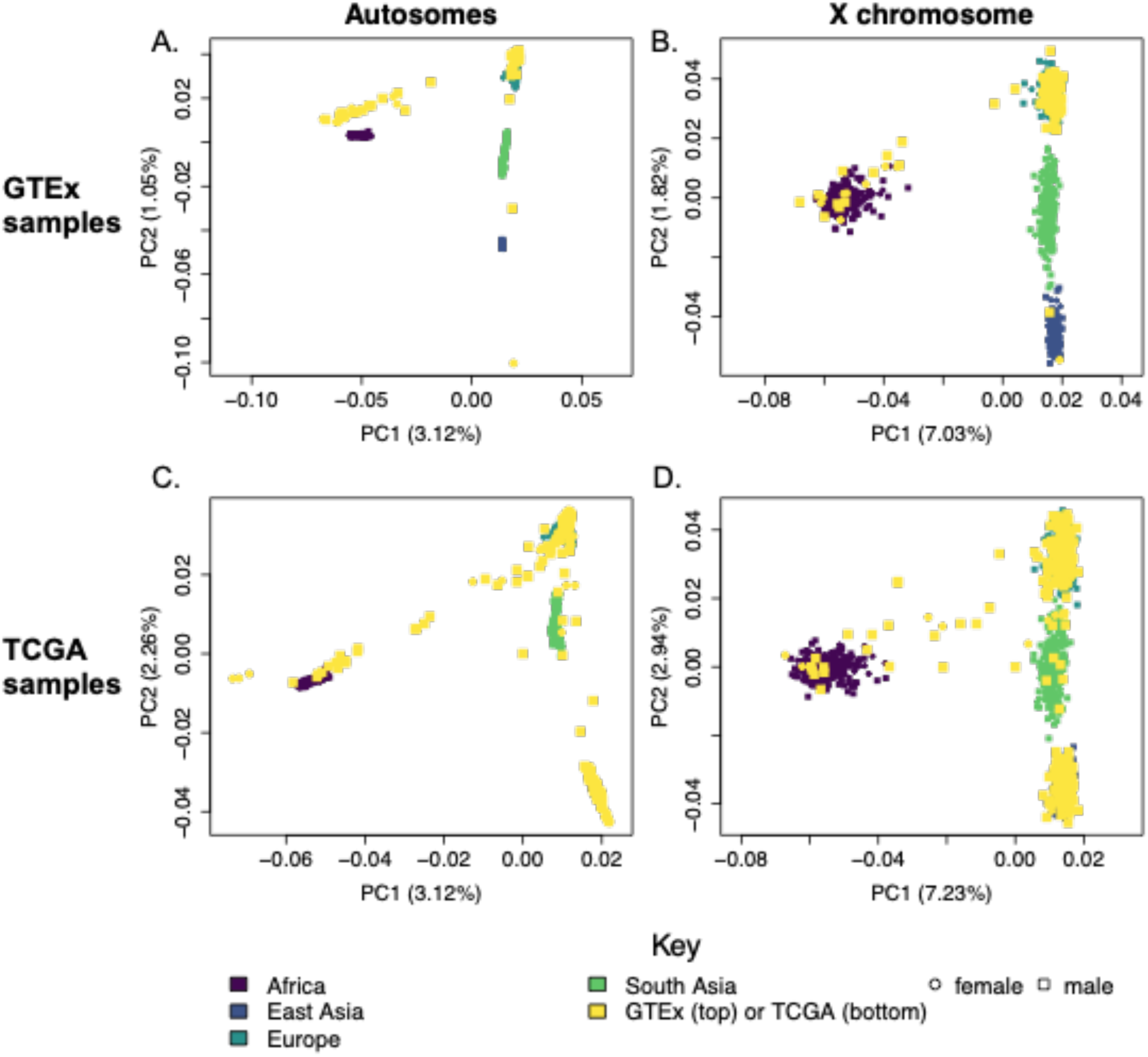
Principal Component Analysis (PCA) output from a sample datasets plotted against the reference dataset. Principal Components 1 and 2 for all individuals for A) autosomes merged and B) X chromosome for the GTEx dataset, and C) autosomes merged and D) X chromosome for the TCGA dataset. Purple points represent the reference samples of African descent, blue points represent reference samples of East Asian descent, dark green points represent reference samples of European descent, and light green represents reference samples of South Asian descent in the 1000 Genomes reference panel. Yellow points represent samples from the sample datasets (GTEx and TCGA).

We find that, using genome-wide genotypes, the genetic ancestry of most study samples does match that which is reported, with notable exceptions, and that we are able to infer ancestry of samples of unreported origin. The inferred ancestry matches closely with the self-reported race information in the GTEx dataset (Supplementary Table 4). One of the GTEx individuals was missing self-reported race. Based on genetic ancestry, this individual was inferred as admixed East Asian and South Asian (Supplementary Table 4). In the TCGA liver cancer dataset, we found 11 individuals with discrepancies between self-reported race and inferred ancestry; for all of these individuals, their self-reported race was white and inferred ancestry was South Asian (Supplementary Table 5). We further inferred ancestry for the 10 individuals in the TCGA liver cancer dataset with no self-reported race (Supplementary Table 5).

We additionally ran PopInf v1.0 on each autosome and the X chromosome separately, finding that chromosome-specific ancestry does not always match that inferred from the whole genome (Figure 2A and C). We identify 16 individuals in the GTEx dataset and 56 individuals in the TCGA dataset (Figure 2 B and D) with variation in chromosome-specific ancestry. All of the admixed individuals had different inferred ancestry results among their chromosomes, as expected. However, there were also 60 (12 from GTEx and 48 from TCGA) individuals inferred as having only one ancestry when analyzing all autosomes together that showed variation in chromosome-specific ancestry (Figure 2 B and D). These ancestry differences across the genome shows that assigning ancestry based only on genome-wide genotypes may result in missing clusters of ancestry across any single chromosome, which may lower our ability to identify risk alleles in datasets consisting of samples of diverse and admixed backgrounds.

**Figure 2.**
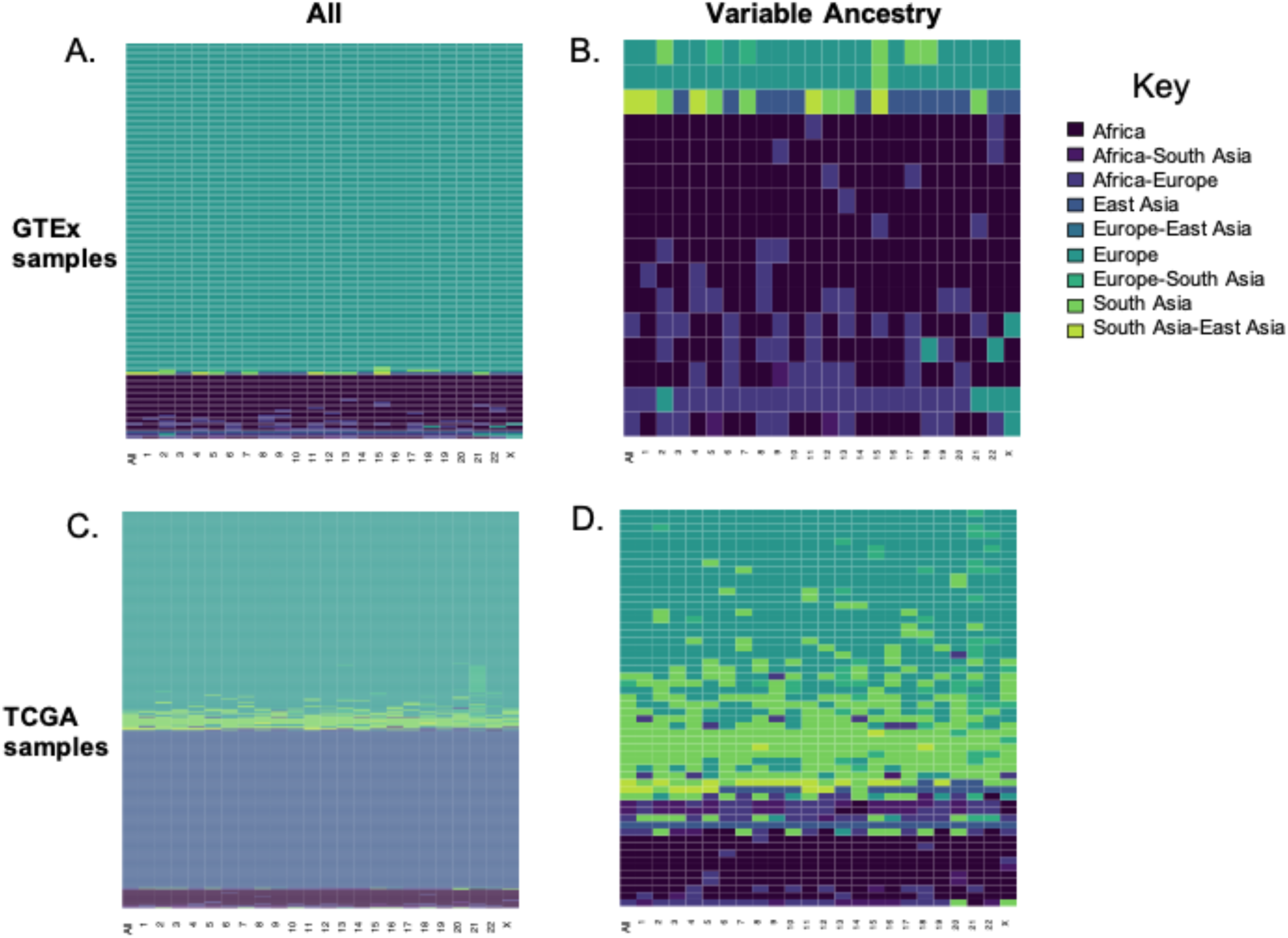
Inferred ancestry for all autosomes combined and each chromosome separately. A) All 148 GTEx individuals, B) the subset of GTEx individuals with variation in inferred ancestry among their chromosomes. C) All 403 TCGA individuals D) the subset of TCGA individuals with variation in inferred ancestry among their chromosomes. Males and females were run together, and only the autosomes and X chromosome were analyzed. The x-axis represents the chromosome analyzed and the y-axis represents the individual from the GTEx dataset. Colors represent inferred ancestry.

## CONCLUSION

Here, we provide a workflow that will set up and run PCA, summarize the PCA output, and provide the user with plots and an easily searchable inferred ancestry report for samples with unknown or unverified population information. Inferred ancestry results from the GTEx and TCGA datasets revealed heterogeneity in ancestry across the genome, and by chromosome. PopInf can be modified to work with any reference panel, and may be applied to similarly infer chromosomal and genome-wide ancestry in diverse populations.

## Acknowledgements

This publication was supported by the National Institute of General Medical Sciences of the National Institutes of Health under Award Number R35GM124827 to MAW. The content is solely the responsibility of the author and does not necessarily represent the official views of the National Institutes of Health. HMN was supported by an ASU Center for Evolution and Medicine postdoctoral fellowship and the Marcia and Frank Carlucci Charitable Foundation postdoctoral award from the Prevent Cancer Foundation. The authors acknowledge Research Computing at Arizona State University for providing high performance computing resources that have contributed to the research results reported within this paper.

## Authors’ contributions

MAW and AMTO conceived the ideas and designed methodology; AMTO and AJD collected the data and analyzed the data. HMN contributed to processing the TCGA data. AMTO and MAW led the writing of the first draft of the manuscript. All authors contributed critically to writing and editing the drafts and gave final approval for publication.

## Data accessibility

PopInf v1.0, processed 1000 Genomes reference file used in this manuscript, and an accompanying tutorial are available on Github: https://github.com/SexChrLab/PopInf.

**Supplementary Table 1. GTEx samples used as the unknown sample dataset.** Here we analyzed the population ancestry from whole genome sequence data from 148 samples available from the GTEx dataset. GTEx (release V6p) whole genome sequence data (dbGaP accession #8834) were downloaded from dbGaP.

**Supplementary Table 2. TCGA samples used as the unknown sample dataset.** Here we analyzed the population ancestry from whole exome sequence data from 403 samples available from the TCGA dataset. TCGA whole exome sequence data (dbGaP accession #11368) were downloaded from NCI Genomic Data Commons (21).

**Supplementary Table 3. 1000 Genomes samples used for this reference panel.** Here we chose 986 unrelated individuals from 1000 Genomes release 3 data downloaded as VCF mapped to GRCh37 from ftp://ftp.1000genomes.ebi.ac.uk/vol1/ftp/release/20130502/. To include global genetic variation in the reference panel, we chose individuals across populations in Africa, Asia, and Europe.

**Supplementary Table 4. GTEx inferred ancestry and self-reported race comparison.** We ran PopInf for all autosomes merged and the X chromosome separately on each individual in the GTEx dataset. We compared these results to the self-reported race information for each individual.

**Supplementary Table 5. TCGA inferred ancestry and self-reported race comparison.** We ran PopInf for all autosomes merged and the X chromosome separately on each individual in the TCGA dataset. We compared these results to the self-reported race information for each individual.

## Notes

https://github.com/SexChrLab/PopInf

## REFERENCES

1. Timpson NJ, Greenwood CMT, Soranzo N, Lawson DJ, Richards JB. Genetic architecture: the shape of the genetic contribution to human traits and disease. Nature Reviews Genetics. 2018;19:110–24.

2. Hindorff LA, Gillanders EM, Manolio TA. Genetic architecture of cancer and other complex diseases: lessons learned and future directions. Carcinogenesis. 2011;32:945–54.

3. Price AL, Zaitlen NA, Reich D, Patterson N. New approaches to population stratification in genome-wide association studies. Nature Reviews Genetics. 2010;11:459–63.

4. Yuan J, Hu Z, Mahal BA, Zhao SD, Kensler KH, Pi J, et al. Integrated Analysis of Genetic Ancestry and Genomic Alterations across Cancers. Cancer Cell. 2018;34:549-560.e9.

5. Dutil J, Chen Z, Monteiro AN, Teer JK, Eschrich SA. An Interactive Resource to Probe Genetic Diversity and Estimated Ancestry in Cancer Cell Lines. Cancer Res. 2019;79:1263–73.

6. Patterson N, Price AL, Reich D. Population Structure and Eigenanalysis. PLoS Genetics. 2006;2:e190.

7. Alexander DH, Novembre J, Lange K. Fast model-based estimation of ancestry in unrelated individuals. Genome Res [Internet]. 2009 [cited 2018 Jul 12]; Available from: http://genome.cshlp.org/content/early/2009/07/31/gr.094052.109

8. Maples BK, Gravel S, Kenny EE, Bustamante CD. RFMix: A Discriminative Modeling Approach for Rapid and Robust Local-Ancestry Inference. The American Journal of Human Genetics. 2013;93:278–88.

9. Pedersen BS, Quinlan AR. Who’s Who? Detecting and Resolving Sample Anomalies in Human DNA Sequencing Studies with Peddy. The American Journal of Human Genetics. 2017;100:406–13.

10. Lonsdale J, Thomas J, Salvatore M, Phillips R, Lo E, Shad S, et al. The Genotype-Tissue Expression (GTEx) project [Internet]. Nature Genetics. 2013 [cited 2018 Jul 5]. Available from: http://www.nature.com/articles/ng.2653

11. Ally A, Balasundaram M, Carlsen R, Chuah E, Clarke A, Dhalla N, et al. Comprehensive and Integrative Genomic Characterization of Hepatocellular Carcinoma. Cell. 2017;169:1327-1341.e23.

12. McKenna A, Hanna M, Banks E, Sivachenko A, Cibulskis K, Kernytsky A, et al. The Genome Analysis Toolkit: A MapReduce framework for analyzing next-generation DNA sequencing data. Genome Res. 2010;20:1297–303.

13. Danecek P, Auton A, Abecasis G, Albers CA, Banks E, DePristo MA, et al. The variant call format and VCFtools. Bioinformatics. 2011;27:2156–8.

14. Quinlan AR, Hall IM. BEDTools: a flexible suite of utilities for comparing genomic features. Bioinformatics. 2010;26:841–2.

15. Chang CC, Chow CC, Tellier LC, Vattikuti S, Purcell SM, Lee JJ. Second-generation PLINK: rising to the challenge of larger and richer datasets. Gigascience. 2015;4:7.

16. R Development Core Team R. R: A language and environment for statistical computing. R foundation for statistical computing Vienna, Austria; 2011.

17. Koster J, Rahmann S. Snakemake--a scalable bioinformatics workflow engine. Bioinformatics. 2012;28:2520–2.

18. Ross MT, Grafham DV, Coffey AJ, Scherer S, McLay K, Muzny D, et al. The DNA sequence of the human X chromosome. Nature. 2005;434:325–37.

19. Skaletsky H, Kuroda-Kawaguchi T, Minx PJ, Cordum HS, Hillier L, Brown LG, et al. The male-specific region of the human Y chromosome is a mosaic of discrete sequence classes. Nature. 2003;423:825–37.

20. Consortium T 1000 GP. A global reference for human genetic variation. Nature. 2015;526:68–74.

21. Grossman RL, Heath AP, Ferretti V, Varmus HE, Lowy DR, Kibbe WA, et al. Toward a Shared Vision for Cancer Genomic Data. New England Journal of Medicine. 2016;375:1109–12.

